# Consumption of Cattle Hide Singed with Scrap Tyre is Detrimental to the Liver, Heart and Kidney of Rats

**DOI:** 10.1101/2020.03.18.996397

**Authors:** Eze. C. Woko, C.O. Ibegbulem, Chinwe. S. Alisi

**Author notes:** Corresponding Author: **Name:** Woko, Eze Chidozie, **Address:** Bioresources Development Centre Arochukwu, National Biotechnology Development Agency, P.M.B 5118 Abuja, Nigeria., **E-mail:**.

## Abstract

The presence of essential amino acids in meat makes it a complete protein, this makes meat a highly sort after source of protein in the human diet. The World’s demand for animal-derived protein has been projected to double by 2050. As a result there is a resultant increase in livestock industries to meet this demand. While meat is generally consumed as a source of protein, processed cattle hide popularly known as “*Kanda”* in southeastern Nigeria is consumed as a substitute for meat though it may not necessarily provide the same level of nutritional value with meat. The method of processing this food delicacy (*Kanda*) by singeing with scrap tyre or firewood has opened the door for heavy metal and or polycyclic aromatic hydrocarbon (PAHs) contamination, thereby putting unsuspecting consumers at health risk. This study therefore investigated the effects of consuming scrap tyre-singed cattle hide and firewood singed cattle hide on the kidney, liver and heart of male Wistar rats. Consuming singed cattle hides significantly elevated some serum marker enzymes important in kidney, liver and heart damage i.e alanine and aspartate aminotransferases, alkaline phosphatase, creatine kinase, and lactate dehydrogenase, with the groups that were fed with the hide singed with scrap tyre showing severe elevations. Consuming the singed *Kanda* also significantly decreased the serum concentrations of K^+^, Na^+^, Cl^−^, and significantly increased HCO_3_^−^ and urea concentration. An examination of the organ tissues also revealed serious morphological changes. In conclusion, consuming singed *Kanda* had detrimental effects on the vital organs studied.

## 1. Introduction

The *Kanda* also known as “*Ponmo*” in Southwestern Nigeria and “*Welle*” in southern Ghana is served as food delicacy in several parts of Africa^[1,2]^. Removal of the hair from the hide in southeastern Nigeria is traditionally done by singeing with firewood, or recently with scrap tyres to get rid of the fur, the burnt hide is scraped to remove ash (black soothe) followed by washing with water to give the finished product (*Kanda*). Owing to the fact that tyres contain many potentially harmful substances: natural and synthetic rubber polymers, oil fillers, sulfur and sulfur compounds, phenolic resin, clay, aromatic, naphthenic and paraffinic oil, fabric, petroleum waxes, pigments such as zinc oxide and titanium dioxide, carbon black, fatty acid**s,** inert materials and fibres made from steel, nylon, polyester or rayon^[3–9]^, singeing *Kanda* with scrap tyres may impose the risk of depositing these elements and compounds into the *kanda*, and expose the unsuspecting consumer to health hazards. It is on this premise that this study investigates the effects of consuming scrap tyre and firewood singed cattle hides on the vital organs (kidney, liver and heart) of Wistar rats.

Kidneys are essential to having a healthy body. They are mainly responsible for filtering waste products, excess water, and other impurities out of the blood. They also regulate pH, salt, and potassium levels in the body^[10]^. Thus, the serum concentrations of various electrolytes like chloride, potassium, bicarbonate, sodium could be used to detect the functional (regulatory) status of the kidney. When the integrity of the kidney is compromised due to exposure to harmful substances or chemicals, its functions are affected and normal health is affected. Same is also applicable to the liver and heart. The liver plays significant role in the body as the organ saddled with the responsibility of metabolizing toxic substances that enter the body. The major functions of the liver can be detrimentally altered by liver injury resulting from acute or chronic exposure to toxicants^[11]^. Enzymes that are found mostly in the liver or other major organs of the body could be used to evaluate the integrity of these organs. Alanine aminotransferase (ALT) and aspartate aminotransferase (AST) are present in the high concentration in the liver. They are present in both the mitochondria and cytosol of the hepatocytes. Damage to this tissue may increase plasma ALT and AST activity^[11]^. Since these enzymes are cytoplasmic in nature, upon liver injury the enzyme enter in to the circulatory system due to altered permeability of membrane. Increases in plasma ALT, AST, alkaline phosphatase (ALP), lactate dehydrogenase (LDH), and creatine kinase (CK) are evidence of liver and heart disease.

The heart is a muscular pump that serves the body in two functions: (1) to collect blood from the tissues of the body and pump it to the lungs and (2) to collect blood from the lungs and pump it to all tissues of the body^[12]^. When the heart is exposed to detrimental levels of toxicants, its functions could be severely affected. Some analysis has established a possible link between exposures to heavy metals or metalloids and risk of conditions such as heart disease, even at low doses — and the greater the exposure, the greater the risk^[13]^. Enzymes such as lactate dehydrogenase (LDH), creatine kinase (CK), and none enzyme protein like troponin I could be used to check the health status of the heart^[14–16]^.

## 2. Materials and methods

### 2.1 Procurement and preparation of samples

Freshly slaughtered cattle hides used for this study were purchased from Ogbor Hill Aba slaughter market, Abia state, Nigeria and processed by the authors. The hide was then divided into two groups; A and B. Sample A was processed by singeing with scrap tyre, the resulting soothe was washed off with clean water. Sample B was processed by singeing with firewood and the resulting soothe was washed off with clean water. Both freshly processed hides were then separately chopped into tiny particles and grated respectively using a locally fabricated grater. The grated particles were then dried and re-grated into smaller particles more suitable for rat consumption.

### 2.2 Study design

Weaned male Wistar rats numbering 35 in total and ranging from 8-9 weeks old with an average live weight of 159.26 g was purchased and used for this study. They were kept at room temperature, fed with standard growers mash rat pellets and water for 7 days (acclimatization period) prior to commencement of the experiment. The rats were divided into 7 groups of 5 rats each and fed for 21 study days as follow:

**Group 1** rats were fed with 10 grams of cattle hide singed with scrap tyre, plus 90 grams of standard grower mash rat pellet feed and water daily for 21 days.
**Group 2** rats were fed with 20 grams of cattle hide singed with scrap tyre, plus 80 grams of standard grower mash rat pellet feed and water daily for 21 days.
**Group 3** rats were fed with 40 grams of cattle hide singed with scrap tyre, plus 60 grams of standard grower mash rat pellet feed and water daily for 21 days.
**Group 4** rats were fed with 10 grams of cattle hide singed with firewood, plus 90 grams of standard grower mash rat pellet feed and water daily for 21 days.
**Group 5** rats were fed with 20 grams of cattle hide singed with firewood, plus 80 grams of standard grower mash rat pellet feed and water daily for 21 days.
**Group 6** rats were fed with 40 grams of cattle hide singed with firewood, plus 60 grams of standard grower mash rat pellet feed and water daily for 21 days.
**Group 7 (control)** rats were fed with 100 grams of standard grower mash rat pellet feed and water daily for 21 days.

### 2.3 Collection of blood and organ samples

Blood samples were collected by orbital technique for assay of biochemical parameters using plain bottles. The blood samples were allowed to cloth before it was centrifuged at 250 rpm for 15 minutes and the serum collected and used for the biochemical analysis. For the organ samples, animals were anaesthetized with dichloromethane and the heart, liver, and kidney were excised and perfused in 10% formal saline for histological studies.

### 2.4 determination of biochemical parameters

The serum sodium ion (Na^+^), potassium ion (K^+^), bicarbonate ion (HCO_3_^−^), chloride ion (Cl^−^), urea, bilirubin, aspartate aminotransferase (AST), alanine aminotransferase (ALT), alkaline phosphatase (ALP), Creatine kinase (CK), Lactate dehydrogenase (LDH), and Troponin I (Trop-I) concentrations were measured spectrophotometrically (Pharmacia LKB Ultospec III) using assay kits (Teco Anaheim, CA, USA) while Na^+^/ K^+^ ratio and ALT/AST ratio were simply calculated. The histological analyses of the tissues were done according to the method of Drury^[17]^.

### 2.5 Statistical analysis

Results of groups are calculated as means ± S.D. and subjected to one-way analysis of variance (ANOVA). Least significant difference (LSD) was used for post-hoc analysis using SPSS statistical software package-SPSS version 21 (SPSS Inc. Chicago, IL, USA).

## 3. Results

Table 1 shows the result of serum electrolyte concentrations, serum urea concentration and sodium/potassium ratios for the six study groups and the control. Table 2 shows the serum activity of the liver enzymes, while table 3 shows the serum activity of some heart enzymes and also troponin-I. The histopathology examination of the kidney, liver and heart are shown in plates 1 to 6. Plate 1 (kidney), plate 3 (liver) and plate 5 (heart) represent rat groups (1, 2 and 3) fed with the cattle hide processed by singeing with scrap tyre and the control (group 7), while plate 2 (kidney), plate 4 (liver) and plate 6 (heart) represent rat groups (4, 5 and 6) fed with the cattle hide processed by singeing with firewood and the control (group 7).

**Plate 1:**
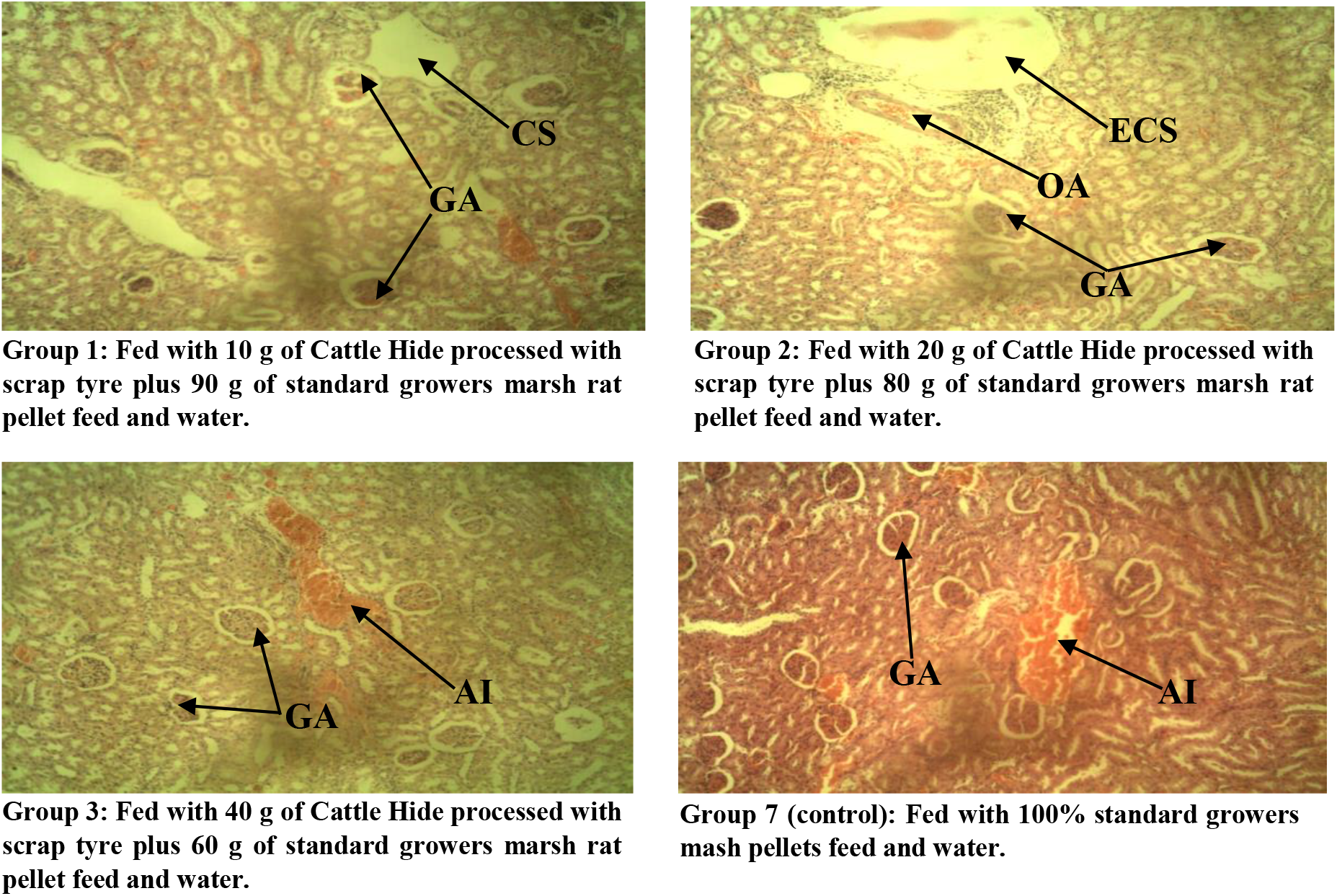
Photomicrograph of the kidney tissues of Wistar rats (male) fed with scrap tyre processed Cattle hide (group 1-3). **Stain H and E.**×100; **CS**= cystic space; **GA**= glomerula atrophy; **ECS** = enlarged cystic space; **OA** = occluded artery; **AI** = acute inflammation.

**Plate 2:**
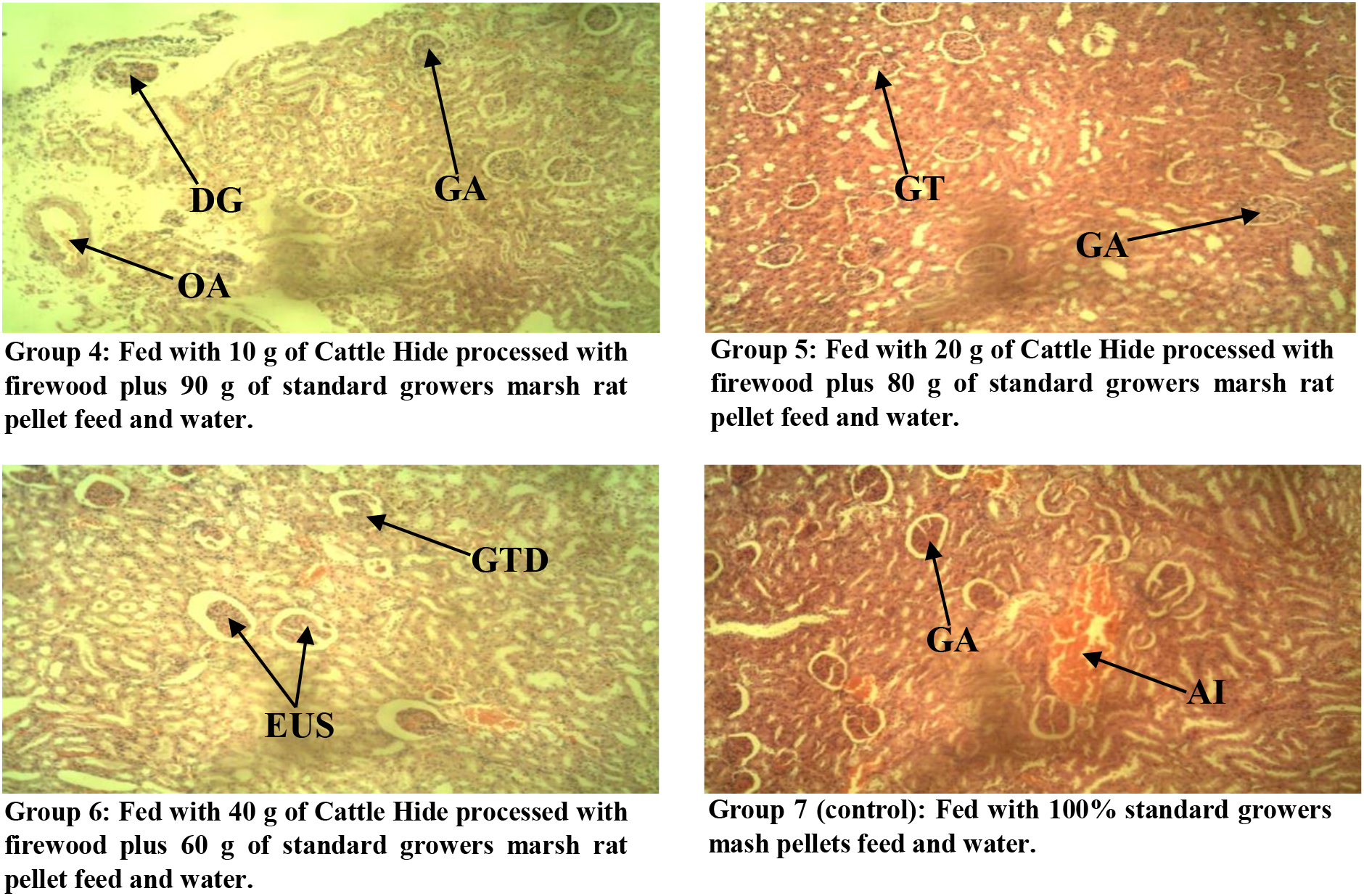
Photomicrograph of the kidney tissues of Wistar rats (male) fed with different concentrations (10 g, 20 g, and 40 g) of the Cattle hide processed with firewood (group 4-6) respectively while group 7 is the control. **Stain: H and E.**×100; **GA** = glomerula atrophy; **DG** = damaged glomeruli; **OA** = occuluded artery; **GT** = glomerula tuft; **GTD** = glomerula tuft degeneration; **EUS** = enlarged urinary space; **AI** = acute inflammation.

**Plate 3:**
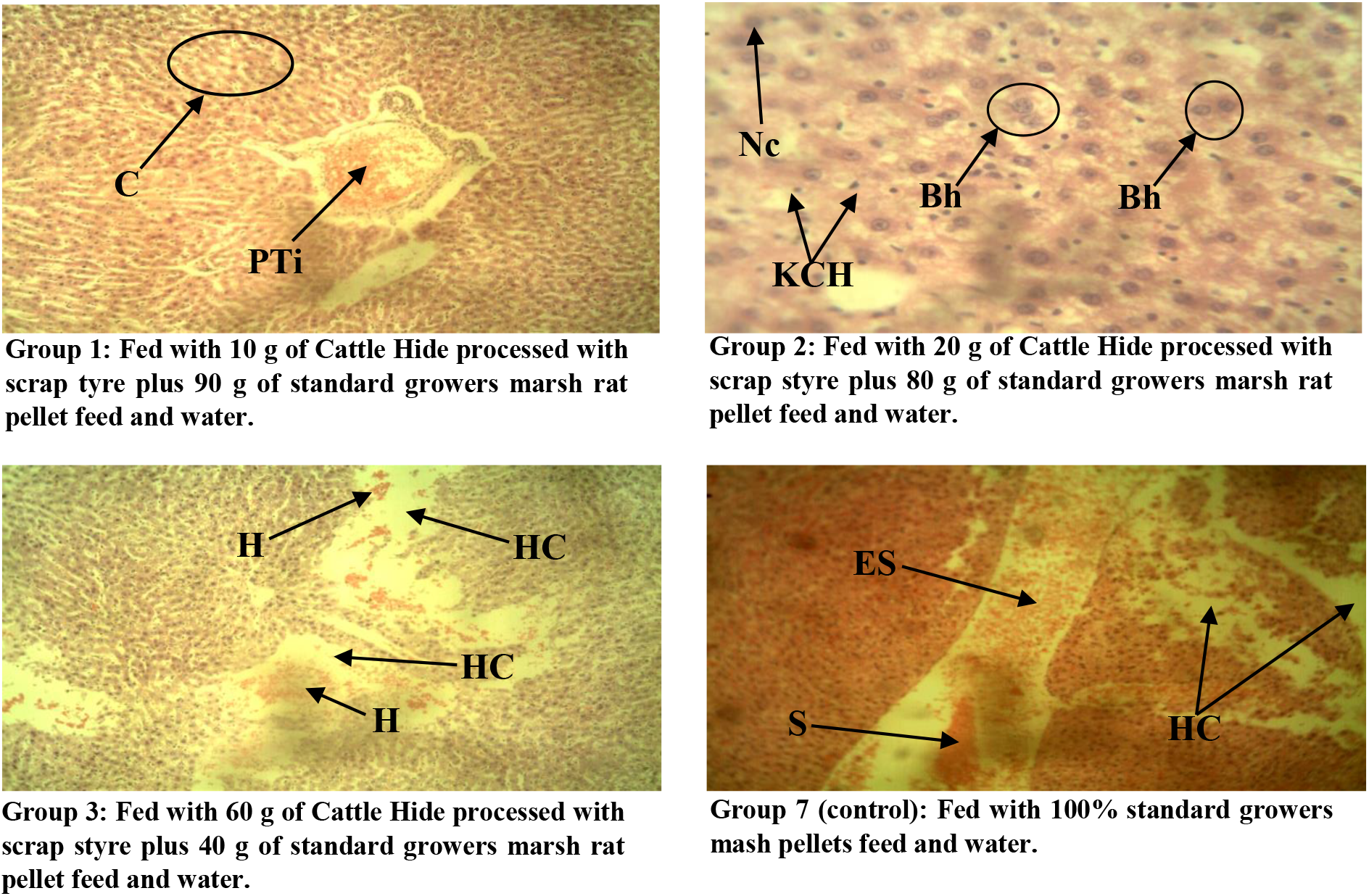
Photomicrograph of the liver tissues of Wistar rats (male) fed with different concentrations (10 g, 20 g, and 40 g) of the Cattle hide processed with scrap tyre, while group 7 is the control; **Stain: H and E.**×100; **C** = Cytoplasmic vacoulation; **PTi** = Portal triad inflammation; **Nc** = Necrotic cells; **Bh** = Binucleation of hepatocytes; **KCH** = Kupfer cells hyperplasia; **H** = Hyperemia; **HC** = distortion of hepatic chord; **S** = Congestion of sinusoid; **ES** = enlarged sinusoid.

**Plate 4:**
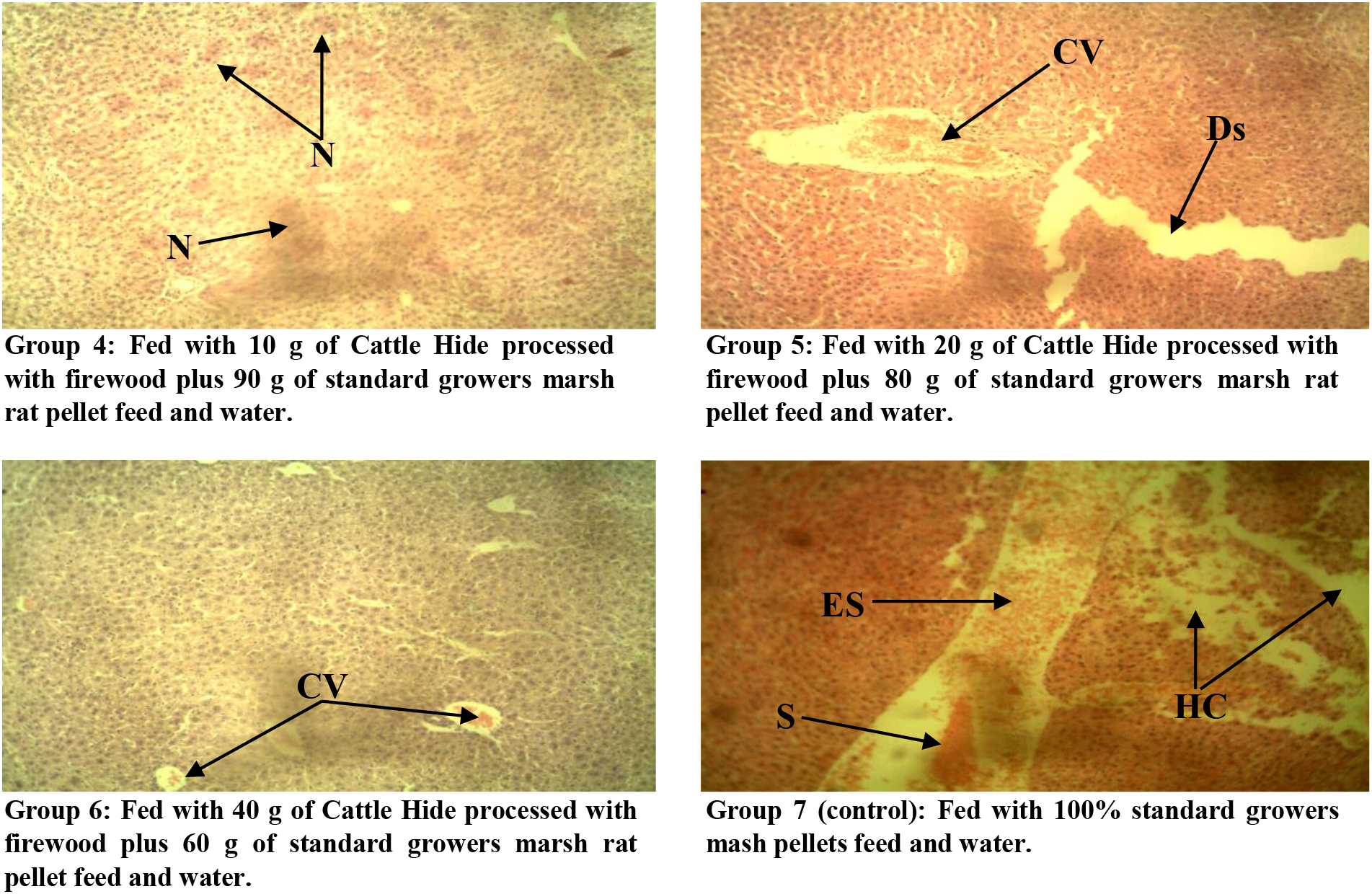
Photomicrograph of the liver tissues of Wistar rats (male) fed with different concentrations (10 g, 20 g, and 40 g) of the Cattle hide processed with firewood (group 4-6) respectively while group 7 is the control. **Stain: H and E.**×100; **N** = multi focal necrosis; **CV** = congested central vein; **DS** = dilation of sinusoid; **HC** = distortion of hepatic chord; **S** = Congestion of sinusoid; **ES** = enlarged sinusoid.

**Plate 5:**
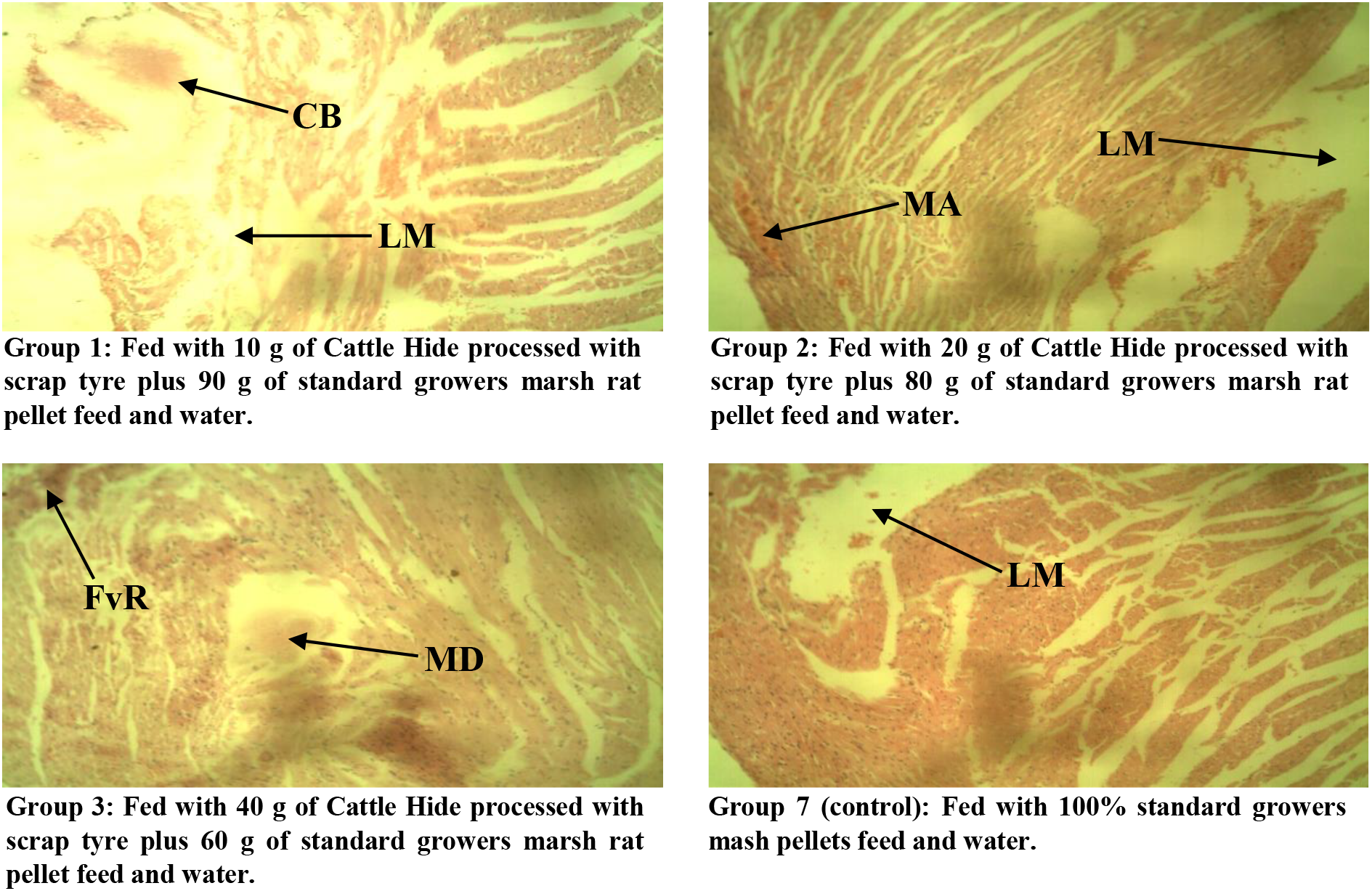
Photomicrograph of the heart tissues of Wistar rats (male) fed with different concentrations (10 g, 20 g, and 40 g) of the Cattle hide processed scrap tyre (group 1-3) respectively. **Stain: H and E**. ×100; **CB** = congested blood; **LM** = loss of myofibres; **MA** = minor arthritis; **FvR** = fibrovascular response; **MD** = myocardial degeneration.

**Plate 6:**
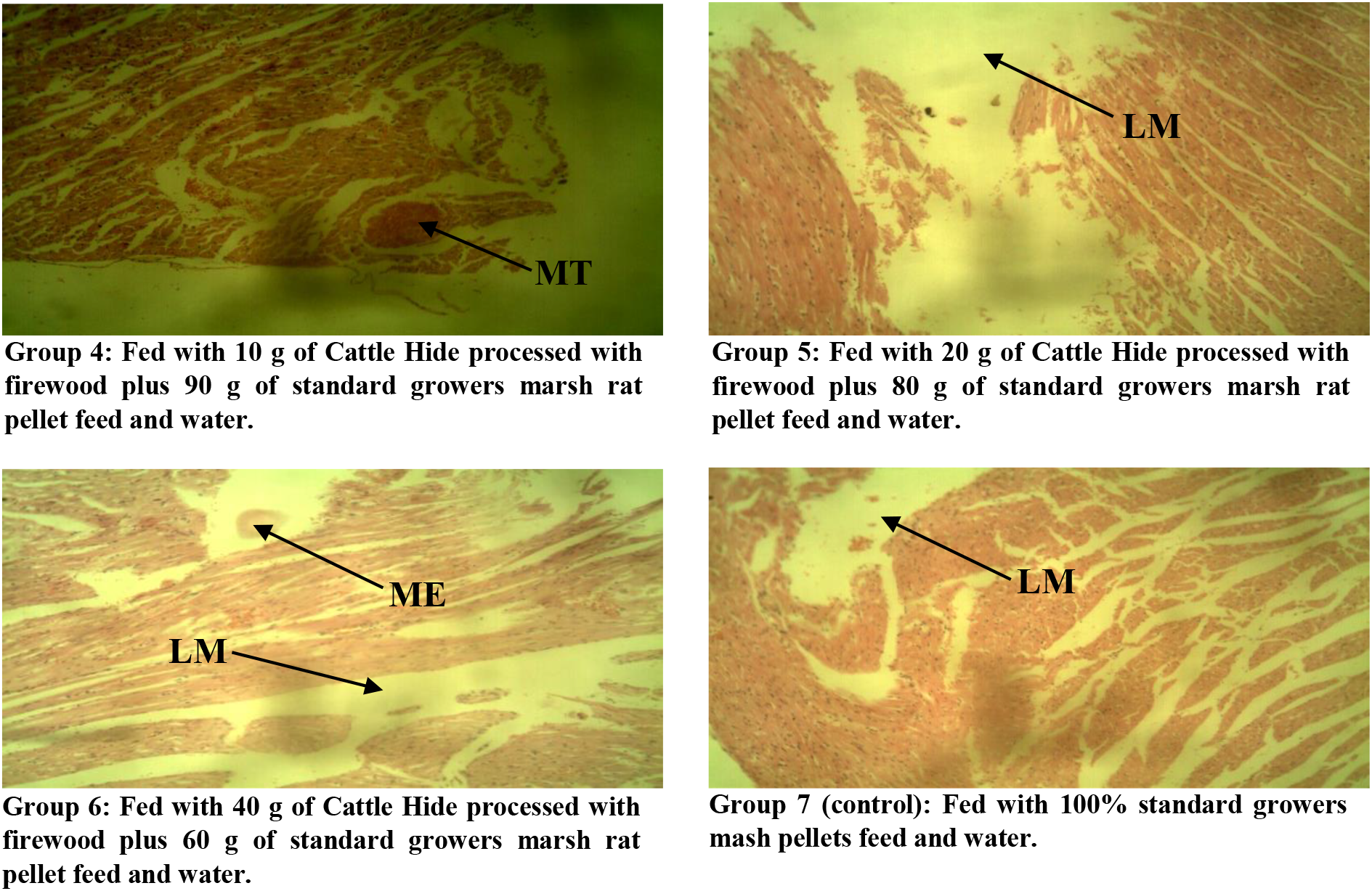
Photomicrograph of the heart tissues of Wistar rats (male) fed with different concentrations (10 g, 20 g, and 40 g) of the Cattle hide processed with firewood (group 4-6) respectively while group 7 is the control. **Stain: H and E**. ×100; **MT** = Minor thrombosis; **ME** = minor edema; **LM** = loss of myofibres.

**Table 1:**
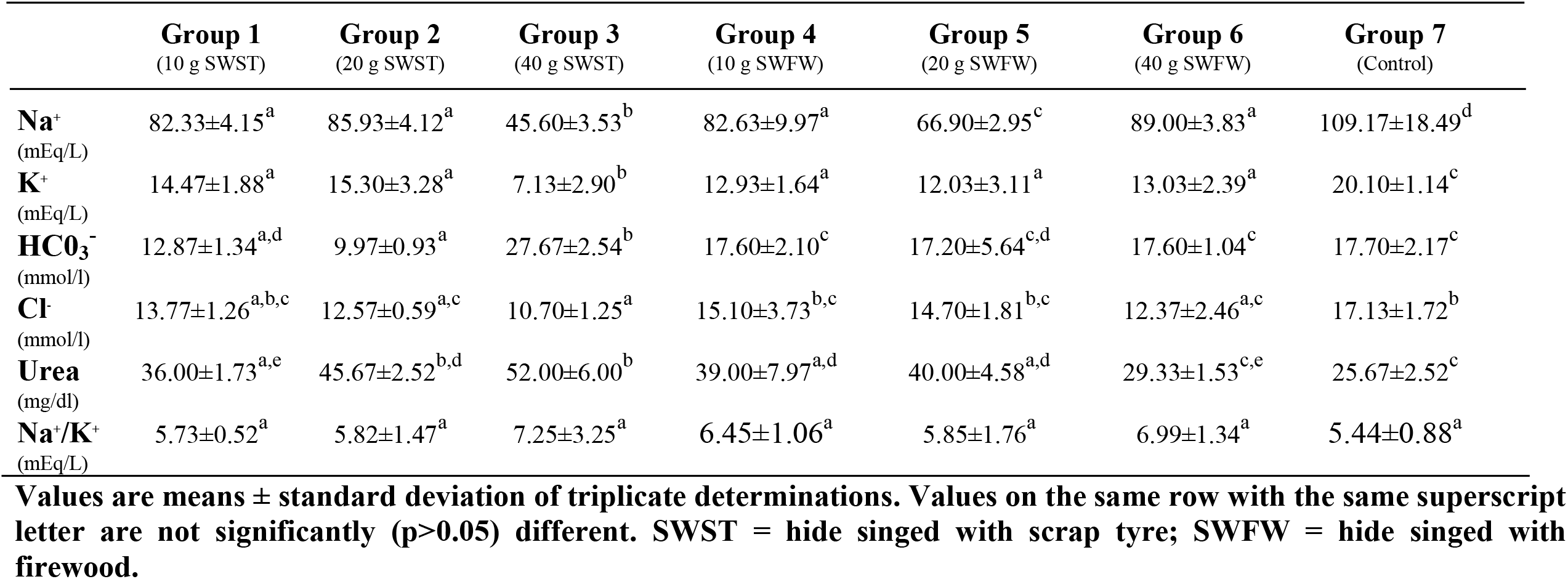
Serum electrolyte/ kidney function test.

|  | **Group 1**<br>(10 g SWST) | **Group 2**<br>(20 g SWST) | **Group 3**<br>(40 g SWST) | **Group 4**<br>(10 g SWFW) | **Group 5**<br>(20 g SWFW) | **Group 6**<br>(40 g SWFW) | **Group 7**<br>(Control) |
| --- | --- | --- | --- | --- | --- | --- | --- |
| **Na<sup>+</sup>**<br>(mEq/L) | 82.33±4.15<sup>a</sup> | 85.93±4.12<sup>a</sup> | 45.60±3.53<sup>b</sup> | 82.63±9.97<sup>a</sup> | 66.90±2.95<sup>c</sup> | 89.00±3.83<sup>a</sup> | 109.17±18.49<sup>d</sup> |
| **K<sup>+</sup>**<br>(mEq/L) | 14.47±1.88<sup>a</sup> | 15.30±3.28<sup>a</sup> | 7.13±2.90<sup>b</sup> | 12.93±1.64<sup>a</sup> | 12.03±3.11<sup>a</sup> | 13.03±2.39<sup>a</sup> | 20.10±1.14<sup>c</sup> |
| **HCO<sub>3</sub><sup>-</sup>**<br>(mmol/l) | 12.87±1.34<sup>a,d</sup> | 9.97±0.93<sup>a</sup> | 27.67±2.54<sup>b</sup> | 17.60±2.10<sup>c</sup> | 17.20±5.64<sup>c,d</sup> | 17.60±1.04<sup>c</sup> | 17.70±2.17<sup>c</sup> |
| **Cl<sup>-</sup>**<br>(mmol/l) | 13.77±1.26<sup>a,b,c</sup> | 12.57±0.59<sup>a,c</sup> | 10.70±1.25<sup>a</sup> | 15.10±3.73<sup>b,c</sup> | 14.70±1.81<sup>b,c</sup> | 12.37±2.46<sup>a,c</sup> | 17.13±1.72<sup>b</sup> |
| **Urea**<br>(mg/dl) | 36.00±1.73<sup>a,e</sup> | 45.67±2.52<sup>b,d</sup> | 52.00±6.00<sup>b</sup> | 39.00±7.97<sup>a,d</sup> | 40.00±4.58<sup>a,d</sup> | 29.33±1.53<sup>c,e</sup> | 25.67±2.52<sup>c</sup> |
| **Na<sup>+</sup>/K<sup>+</sup>**<br>(mEq/L) | 5.73±0.52<sup>a</sup> | 5.82±1.47<sup>a</sup> | 7.25±3.25<sup>a</sup> | 6.45±1.06<sup>a</sup> | 5.85±1.76<sup>a</sup> | 6.99±1.34<sup>a</sup> | 5.44±0.88<sup>a</sup> |

**Values are means ± standard deviation of triplicate determinations. Values on the same row with the same superscript letter are not significantly (p>0.05) different. SWST = hide singed with scrap tyre; SWFW = hide singed with firewood.**

**Table 2:**
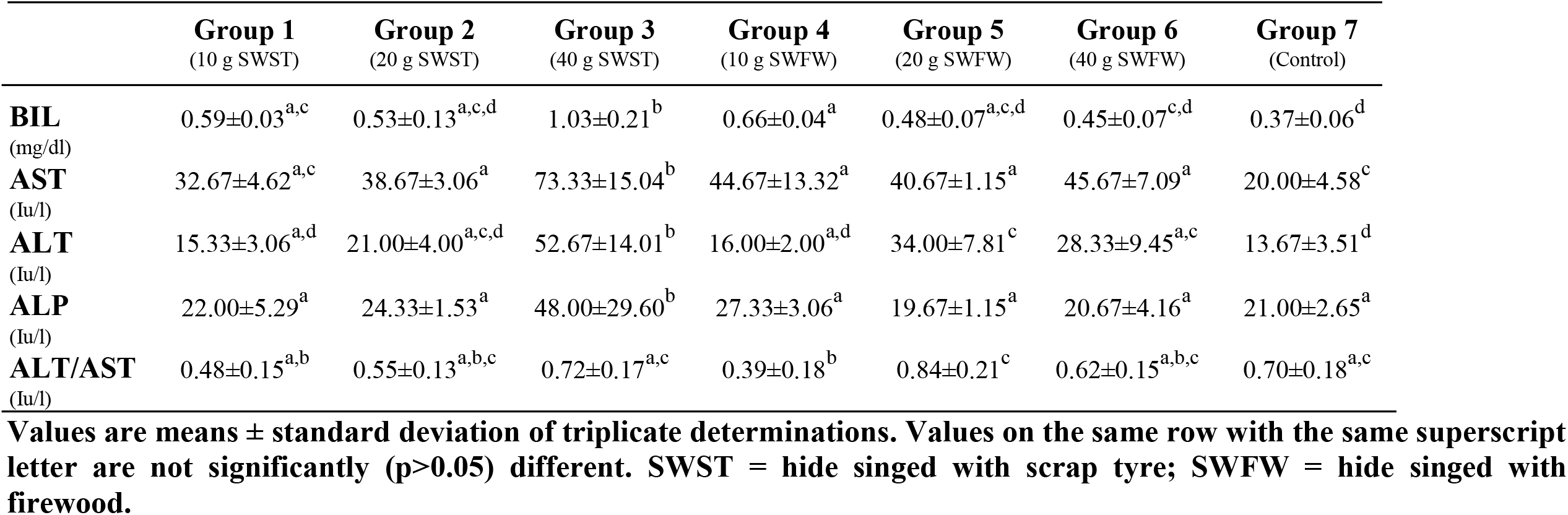
Liver function test result.

**Table 3:** Heart function test.

|  | **Group 1**<br>(10 g SWST) | **Group 2**<br>(20 g SWST) | **Group 3**<br>(40 g SWST) | **Group 4**<br>(10 g SWFW) | **Group 5**<br>(20 g SWFW) | **Group 6**<br>(40 g SWFW) | **Group 7**<br>(Control) |
| --- | --- | --- | --- | --- | --- | --- | --- |
| **CK**<br>(u/l) | 71.87±2.21<sup>a</sup> | 100.13±12.43<sup>a</sup> | 221.93±83.41<sup>b</sup> | 59.33±3.42<sup>a</sup> | 74.63±21.20<sup>a</sup> | 68.03±5.89<sup>a</sup> | 54.60±5.26<sup>a</sup> |
| **LDH**<br>(u/l) | 168.77±3.75<sup>a,c,e</sup> | 202.13±35.75<sup>a,b</sup> | 212.43±37.70<sup>b</sup> | 150.63±22.26<sup>c,d,e</sup> | 119.43±20.45<sup>d,e</sup> | 163.17±16.01<sup>a,c,e</sup> | 127.43±6.63<sup>e</sup> |
| **TROP-I**<br>(ng/ml) | 0.80±0.03<sup>a,b</sup> | 0.90±0.08<sup>b</sup> | 1.06±0.12<sup>c</sup> | 0.68±0.04<sup>a,d,e</sup> | 0.63±0.12<sup>d,e</sup> | 0.73±0.06<sup>a,d</sup> | 0.55±0.09<sup>e</sup> |

Values are means ± standard deviation of triplicate determinations. Values on the same row with the same superscript letter are not significantly (p>0.05) different. SWST = hide singed with scrap tyre; SWFW = hide singed with firewood.

## 4. Discussion

The kidneys play important role in serum electrolyte homeostasis. The concentrations of various serum electrolytes could therefore be used to access the severity of kidney damage. With progressive loss of kidney function there is an inevitable occurrence of derangements in electrolytes and acid-base^[18]^. In all the study groups (1-6) low serum sodium (hyponatraemia), low serum potassium (hypokalemia), and low serum chloride (hypochloremia) concentrations were observed when compared to the control, these significant (p>0.05) decreases is evidence of kidney dysfunction in the study groups. As a result of kidney dysfunction sodium is not reabsorbed by the renal tubules and is thus excreted in the urine frequently resulting in hyponatraemia; there is also a progressive failure to excrete water^[19]^. Sodium is essential in the human body; it has a vital role in maintaining the concentration and volume of the extracellular fluid and accounts for most of the osmotic activity of the plasma. Its serum levels are maintained by feedback loops involving the kidney, adrenal glands and hypothalamus. When the serum sodium is low as observed in the study groups (usually because total body water is high), antidiuretic hormone (ADH) is suppressed and dilute urine is excreted. In addition, the kidney produces rennin which stimulates aldosterone production that decreases the excretion of sodium in the urine, therefore increasing sodium levels in the body. It could be therefore that the rats in the study groups due to kidney failure or damage could not among other things produce rennin which in turn caused sodium to be excreted in the urine without reabsorption, a condition that resulted to the low serum sodium (hyponatraemia) as observed in all the study groups. Group 3 had the most critical condition of hyponatraemia, hypokalemia and hypochloremia, this suggests that consuming scrap tyre-singed cattle hide at a higher concentration poses greater health risk. Hypokalemia is a metabolic imbalance characterized by extremely low potassium levels in the blood^[20]^. Under normal circumstances, the kidneys are responsible for excreting 90% of the potassium that is consumed daily with remaining 10% excreted in the feces. Abnormalities of serum potassium are common in patients with chronic kidney disease (CKD), although hyperkalemia (high serum potassium) is a well recognized complication of CKD, the prevalence rates of hyperkalemia (14% - 20%) and hypokalemia (12% - 18%) are similar^[21]^. When kidneys are functioning normally the amount of K^+^ in the diet is sufficient for use by the body and the excess is usually excreted through urine and sweat. When hypokalemia occurs (as observed in the study groups), there is an imbalance resulting from a dysfunction in the normal process or rapid loss of urine or sweat without replacement of sufficient potassium. Hypokalemia has also been reported to impair the ability of the kidneys to concentrate urine, resulting in excessive urination (polyuria) and excessive thirst (polydipsia). The excessive excretion of K^+^ in the urine (kaliuresis) which caused the hypokalemia in the study groups may result from kidney disorders among other factors^[20]^.

According to Zahra, the extremely high bicarbonate and low chloride (hypochloremia) in the study group 3 is almost certainly due to metabolic alkalosis^[22]^. This alkalosis is maintained and potentiated by the kidneys’ inability to excrete excess bicarbonate because of hypochloremia. Hypokalemia is frequently associated with metabolic alkalosis because kaliuresis is increased in response to low H^+^ concentration^[22]^. All the groups fed with the hide singed with firewood (4-6) showed no significant (p<0.05) difference compared to the control while the groups (1-2) fed with the hide singed with scrap tyre showed significant (p<0.05) decrease in serum bicarbonate concentration except group 3 which showed significant (p<0.05) increase compared to the control and other study groups. Elevated serum bicarbonate concentration has been associated with heart failure exacerbation and mortality in patients with chronic kidney disease in an alarming rate^[23]^. Metabolic acidosis leads to chronic kidney disease (CKD); in elderly CKD patients, serum bicarbonate levels is independently associated with CKD progression, and a high serum bicarbonate level is associated with a low risk of CKD progression provided it is within range^[24]^. There was an observed significant (p<0.05) decrease in serum chloride concentrations of study groups 2, 3 and 6 when compared to the control. Chloride is the major anion associated with sodium in ECF. The kidneys are responsible for the maintenance of total body chloride balance. They maintain homeostasis because each kidney is composed of 1 million functional units, nephrons. Part or all of the Cl^−^ filtered by the initial portion of each of the nephron, the glomerulus, will be reabsorbed as a result of both active and passive transport processes along the tubules comprising each nephron. The ability of the nephrons to reabsorb chloride maintains the serum (and ECF) chloride concentration within a narrow range^[25]^. The inability of the kidney to reabsorb the chloride led to the hypochloremia observed in groups 2, 3, and 6. This could be as a result of over-hydration and or congestive heart failure where chloride is retained with sodium retention but is diluted by excess total body water.

Except group 6, all the study groups when compared to the control showed significant (p<0.05) increase in their serum urea concentrations with group 3 having the highest concentration. Urea is a waste product of protein metabolism and digestion formed in the liver and excreted by the kidneys in urine. Therefore urea is directly related to the metabolic function of the liver and the excretory function of the kidney. Kidney disease has been associated with reduced urea excretion and consequent rise in blood concentration as observed in the study groups^[26]^. In fact, nearly all renal diseases cause an inadequate excretion of urea, which causes the blood concentration to rise above normal^[27–30]^. Groups 1, 2, 3, 4 and 5 had significantly increased (p<0.05) urea concentration compared to the control. There was no significant (p<0.05) difference in the sodium-potassium (Na^+^-K^+^) ratio of all the study groups compared to the control or among themselves.

The liver has long been considered the major target organ for most of the chemicals implicated in eliciting toxic effects following environmental exposure^[31]^. All the study groups except 2, 5 and 6 showed significant (p<0.05) increase in the serum bilirubin concentration when compared to the control. Serum bilirubin is an endogenous anion derived from hemoglobin degradation from the red blood cell, abnormal levels in the serum is an indication of underlying liver disease as observed in study groups 1, 3 and 4^[32]^. The observed increment in the activities of AST in serum of group 2, 3, 4, 5, and 6 may be mainly due to the leakage of these enzymes from the liver cytosol into the blood stream as a result of liver damage (necrosis as shown in the histopathology study of the liver tissues). Study groups 3, 5, and 6 showed significant (p<0.05) increase in their serum alanine aminotranferase (ALT) when compared to the control. ALT is found predominately in the liver; lesser quantities are found in the kidneys, heart, and skeletal muscle. Injury or disease affecting the liver functioning part of the organ (parenchyma) will cause a release of this hepatocellular enzyme into the bloodstream, therefore elevating the ALT serum levels. Most ALT serum level increases are due to liver dysfunction. ALT serum levels are quite specific for hepatocellular disease indicators, mildly increased levels results to myocardial infarction, moderately increased level results to hepatic tumor while significantly increased levels result to hepatic necrosis as observed in the liver tissue examination of groups 2 and 4 rats^[30]^. The ALT/AST ratios of only study groups 1, and 4 were significantly (p<0.05) low compared to the control. In viral hepatitis the ALT/AST ratio is greater than 1, while in hepatocellular disease the ratio is less than 1 u/L^[30]^. Group 3 rats are the only group with a significant (p<0.05) increase in alkaline phosphatase (ALP) and creatine kinase (CK) concentrations. Increased levels of ALP as observed with group 3 may signify primary cirrhosis, primary or metastatic liver tumor, and myocardial infarction among others^[30]^. Study groups 2, and 3 showed significant (p<0.05) increase in the serum lactate dehydrogenase (LDH) concentration compared to the control while groups 1, 4, 5 and 6 showed no significant (p<0.05) difference compared to the control. Groups 1, 2, 3 and 6 showed significant increase (p<0.05) in their Troponin-I concentration when compared to the control. Cardiac troponins are the biochemical markers for cardiac disease^[30]^. Cardiac troponins become elevated sooner and remain elevated longer than CK. This expands the time window of opportunity for diagnosis and thrombolytic treatment of myocardial injury. Any rise in creatine levels is only observed if there is marked damage to functional nephrons^[33]^.

Plate 1 shows the photomicrograph of kidney tissues for groups 1 to 3 which were fed with the hide singed with scrap tyre and the control (group 7). In group 1, clear glomerular atrophy was observed with minor cystic space. According to Leh, glomerular atrophy is an acquired diminution of the size of the capillary loops of the kidney associated with wasting as from death and reabsorption of cells, reduced function or malfunction, or hormonal changes^[34]^. Group 2 showed enlarged cystic space with presence of congested blood. The blood vessels especially the artery were also occluded. According to another report, simple kidney cysts are abnormal fluid-filled sacs that form in the kidneys^[35]^. The report further stated that the cause of simple kidney cysts is not fully understood though obstruction of tubules (tiny structures within the kidney that collect urine) or deficiency of blood supply to the kidneys may play a role^[35]^. Group 3 showed acute inflammations and glomerular atrophy. While in group 7 (control) the glomeruli were intact although minor acute inflammation was observed.

Plate 2 shows the photomicrograph of kidney tissues for groups 4 to 6 fed with the hide singed with firewood and the control (group 7). In group 4 the glomeruli were damaged with loss of glomerular contents, there was also the presence of arteries with minor occlusion. Renal artery occlusion (RAO) is a rare but significant cause of kidney loss which may be caused by renal artery thrombosis^[36]^. Group 5 showed minor atrophy of the glomerular tuft with slight distention of the bowman’s space. In group 6 atrophy of glomerular tuft appeared more severe with enlarged urinary space. While in group 7 (control) the glomeruli were intact although minor acute inflammation was observed.

Plate 3 showed the photomicrograph of liver tissues for groups 1, 2 and 3 fed with the hide singed with scrap tyre and the control (group 7). In group 1, portal triad inflammation was observed with cytoplasmic vacuolations. Cytoplasmic vacuolation is a well-known morphological phenomenon observed in mammalian cells after exposure to bacterial or viral pathogens as well as to various natural and artificial low molecular weight compounds. Vacuolation often accompany cell death; however, its role in cell death process remains unclear^[37]^. In the liver, as in every vascularized tissue, cell death (necrosis) evokes an inflammatory reaction which is morphologically manifested by the appearance of inflammatory cells together with edema and congestion around parenchymal cells which in the case of the liver are damaged hepatocytes or damaged bile ducts or damaged blood vessels. Portal inflammation may involve all or some portal fields. In group 2, Kupfer cell hyperplasia, binucleation (double nuclei) of hepatocytes and necrosis of the hepatocytes was observed. Kupfer cells reside within the lumen of the liver sinusoids; therefore they are the first to be exposed to materials absorbed from the gastrointestinal tract^[38]^. Hepatocytes are indeed the major cell type in the liver, they might represent primary modulators of hepatic immunity, especially in the setting of chronic liver injury, where non-injured hepatocytes in close proximity are predisposed to inflammatory mediators as bystanders^[38]^. Group 3 showed distortion of the hepatic chords with loss of hepatocytes, and hyperemia (the buildup of blood). Distortion of hepatic chords is certainly a sign of cirrhosis^[38]^. Cirrhosis is a consequence of chronic liver disease characterized by replacement of liver tissue by dense fibrous scar tissue as well as regenerative nodules formation which result in widespread distortion of normal hepatic architecture^[38]^. The control (group 7) showed congestion, and enlargement of the sinusoid, with distortion of hepatic cord. The reason for these occurrences in the control is inexplicable.

Plate 4 showed photomicrograph of liver tissues for groups 4, 5, and 6 fed with the hide singed with firewood and group 7 (control). In group 4, multifocal necrosis was observed. Group 5 showed enlargement of the sinusoids and congestion of the central vein. In group 6 the central veins were also congested, while group 7 (control) showed congestion, and enlargement of the sinusoid, with distortion of hepatic cord.

Plate 5 shows the photomicrograph of the heart tissues for groups 1 to 3 fed with the hide singed with scrap tyre and the control (group 7). In Group 1, there was typical cardiac muscle (myocardium), blood congestion and degeneration of the myofibres which led to myonecrosis. In Group 2, minor arteritis appeared due to inflammatory infilterates of the lymphocytes, and neutrophils which led to narrowing and obliteration of the lumen which is still at the early stage of arteritis. Loss of myofiber was also observed. Group 3 showed fibrovascular responses with prominent granulation tissues and myocardial degeneration due to circulatory disturbances especially infarction which creates hypoxia and death of myofibers. In Group 7, there was edema and the presence of diffused degenerated myofibers which are observable conditions that led to cardiomyopathy. The above condition showed progressive murine cardiomyopathy, and why these occurred in the control is inexplicable

Plate 6 shows the photomicrograph of the heart tissues for groups 4 to 6 fed with the hide singed with firewood and the control (group 7). In group 4, there appeared the presence of minor thrombosis with typical endocardial myxomatous change usually consisting of focal or diffuse thickening due to presence of myxomatous tissue. In Groups 5 and 6, there was an observed degeneration of the myofibers. In Group 7, there was edema and the presence of diffused degenerated myofibers which are observable conditions that led to cardiomyopathy.

## CONCLUSION

The consumption of cattle hides singed with firewood or scrap tyre as sources of fuel poses significant health hazard to the vital organs of the body like kidney, liver and heart with the hide singed with scrap tyre posing the higher level of risk. Consuming these hides could cause such health challenges as hyponatraemia, hypokalemia, hypochloremia, chronic kidney disease (CKD), cirrhosis, congestive heart failure etc. With the findings of this study, it is advised that eating of scrap tyre or firewood singed cattle hides (*Kanda, Ponmo*, or *Welle*) should be discouraged.

## Presentation at a meeting

The authors declare that this work has not been presented in any meeting.

## Contribution Details

|  | **Woko, E.C** | **Ibegbulem, C.O** | **Alisi, C.S** |
| --- | --- | --- | --- |
| Concepts | Yes | Yes | Yes |
| Design | Yes | Yes | Yes |
| Definition of intellectual content | Yes | Yes | Yes |
| Literature search | Yes | No | No |
| Clinical studies |  |  |  |
| Experimental studies | Yes | Yes | Yes |
| Data acquisition | Yes | Yes | Yes |
| Data analysis | Yes | Yes | Yes |
| Statistical analysis | Yes | Yes | Yes |
| Manuscript preparation | Yes | No | No |
| Manuscript editing | Yes | Yes | Yes |
| Manuscript review | Yes | Yes | Yes |
| Guarantor |  |  |  |

## KEY MESSAGES

There is a danger of contaminating hides when they are singed with tyre or firewood These contamination makes the hide unhealthy for consumption and has been shown by this study to cause damages to the severity of the heart, liver and kidney of Wistar rats studied.

